# *Vibrio cholerae* lineage and pangenome diversity varies geographically across Bangladesh over one year

**DOI:** 10.1101/2024.11.12.623281

**Authors:** Chuhan Qin, Patrick Lypaczewski, Md. Abu Sayeed, Aline C. Cuénod, Lindsey Brinkley, Ashton Creasy-Marrazzo, Emilee T. Cato, Kamrul Islam, Md. Imam Ul Khabir, Md. Taufiqur R. Bhuiyan, Yasmin Begum, Firdausi Qadri, Ashraful I. Khan, Eric J. Nelson, B. Jesse Shapiro

## Abstract

Cholera is a diarrhoeal disease caused by *Vibrio cholerae*. It remains a major public health challenge in the endemic region around the Bay of Bengal. Over decadal time scales, one lineage typically dominates the others and spreads in global pandemic waves. However, it remains unclear to what extent diverse lineages co-circulate during a single outbreak season. Defining the pool of diversity during finer time scales is important because the selective pressures that impact *V. cholerae* – namely antibiotics and phages – are dynamic on these time scales. To study the nationwide diversity of *V. cholerae*, we long-read sequenced 273 *V. cholerae* genomes from seven hospitals over one year (2018) in Bangladesh. Four major *V. cholerae* lineages were identified: known lineages BD-1, BD-2a, and BD-2b, and a novel lineage that we call BD-3. In 2022, BD-1 caused a large cholera outbreak in Dhaka, apparently outcompeting BD-2 lineages. We show that, in 2018, BD-1 was already dominant in the five northern regions, including Dhaka, consistent with an origin from India in the north. By contrast, we observed a higher diversity of lineages in the two southern regions near the coast. The four lineages differed in pangenome content, including integrative and conjugative elements (ICEs) and genes involved in resistance to bacteriophages and antibiotics. Notably, BD-2a lacked an ICE and is predicted to be more sensitive to phages and antibiotics, but nevertheless persisted throughout the year-long sampling period. Genes associated with antibiotic resistance in *V. cholerae* from Bangladesh in 2006 were entirely absent from all lineages in 2018-19, suggesting shifting costs and benefits of encoding these genes. Together, our results highlight the dynamic nature of the *V. cholerae* pangenome and the geographic structure of its lineage diversity.

## Introduction

Cholera is an acute bacterial infectious disease characterised by profuse watery diarrhoea. It is caused by the Gram-negative pathogen *Vibrio cholerae*, which is found in coastal brackish waters in planktonic forms or attached to zooplankton or shellfish (1,2). *V. cholerae* is genetically diverse in the aquatic environment, and only two serotypes (O1 and O139) are associated with pandemic cholera in humans (2). The current 7^th^ pandemic was first detected in 1961 in Indonesia, caused by a *V. cholerae* lineage referred to as the 7^th^ Pandemic El Tor (7PET), which is genetically distinct from the Classical Type responsible for the 6^th^ pandemic (3). Currently, the public health burden of cholera remains high, with 2.9 million cases and 95,000 deaths estimated to be caused by cholera every year (4). The burden of cholera is likely underestimated, as most cholera infections do not display any symptoms (5). A recent study based on serological surveillance showed that the *V. cholerae* infection rate may be much higher – at 535 per 1,000 people annually in Bangladesh (5). Outside the endemic region around Bangladesh, shorter but often devastating outbreaks can occur, as demonstrated in 2010 in Haiti, or in 2016 in Yemen (3,6).

In 2022, Bangladesh experienced one of the largest cholera outbreaks in recent years when the BD-2 *V. cholerae* lineage, which had been dominant in Bangladesh since the early 2010s, was superseded by BD-1.2 (7,8). The BD-1.2 lineage is most closely related to BD-1 genomes sampled from India, suggesting an Indian origin (7). BD-1 and BD-2 are relatively closely related lineages within the 7PET, but differ by single nucleotide changes and gene content differences in pathogenicity islands such as VSP-II and integrative and conjugative elements (ICEs). It remains unclear which, if any, of these differences explain why one lineage might replace another.

ICEs encode numerous antimicrobial resistance (AMR) genes, and were recently shown to confer resistance to the lytic ICP1 bacteriophage through diverse phage defence genes (9,10). The first ICE discovered in *V. cholerae* was named SXT, as it conferred resistance to <u>S</u>ulfametho<u>X</u>azole and trimethoprim) (11,12). Different ICE variants encode different AMR and phage resistance genes. For example, in a sample of *V. cholerae* genomes from 2006, the genes *qnr_Vc_* (encoding a pentapeptide that protects DNA gyrase from the attack of ciprofloxacin) and *mphA* (encoding a phosphotransferase that inactivates azithromycin) were respectively associated with ciprofloxacin and azithromycin resistance phenotypes localised to the ICE (13). Frequent homologous recombination between ICEs is thought to occur, allowing rapid gain and loss of AMR genes (14). The same is likely true of phage-resistance genes. Notably, the *Vch*Ind5 ICE variant, but not the *Vc*Ind6 ICE, encodes a BREX anti-phage system that is associated with lower phage to *V. cholerae* ratios within patients (9,10). Some *V. cholerae* genomes lack an ICE entirely, suggesting fitness costs under certain conditions.

To determine the distribution of *V. cholerae* lineages and genes circulating in Bangladesh, we sequenced 273 *V. cholerae* genomes collected in 2018 and early 2019 from seven hospitals across the country, spanning a single annual outbreak season. In addition to the previously described BD-1 and BD-2 lineages, we identified two BD-2 sublineages, one of which typically lacks an ICE, and a novel lineage, BD-3, which was restricted to Dhaka in our data set. Despite the higher prevalence of BD-1, BD-2 persisted during the sampled period. BD-1 was more prevalent in northern Bangladesh bordering India, supporting its Indian origin. Southern regions contained a higher diversity of lineages, potentially due to distinct environments and migration patterns. We also highlight the diversity of AMR and phage resistance genes, many of which occur on the ICE and differ across lineages. Two genes associated with antibiotic resistance in 2006 were absent from all *V. cholerae* genomes in our 2018-19 sample, exemplifying the dynamic nature of the pangenome.

## Methods

### Ethics Statement

Samples for this study were collected as part of two published IRB-approved clinical studies in Bangladesh: (i) The mHealth Diarrhoea Management (mHDM) cluster randomised controlled trial (IEDCR IRB/2017/10; icddr,b ERC/RRC PR-17036; University of Florida IRB 201601762) (15). (ii) The National Cholera Surveillance (NCS) study (icddr,b ERC/RRC PR-15127) (16).

### Study Design

A prospective longitudinal study of patients presenting with diarrhoeal disease was conducted at five Bangladesh Ministry of Health and Family Welfare district hospitals (both mHDM and NCS sites) and two centralised exclusive NCS hospitals (BITID; icddr,b). Sites were distributed geographically nationwide (9). For mHDM, the inclusion criteria were patients with acute uncomplicated diarrhoea (less than 7 days with at least 3 loose stools in the last 24 hours) and age greater than or equal to two months; 4 patients per day were sampled. For NCS, the case definitions differed by age. For age under two months: changed stool habit from the usual pattern in terms of frequency (more than the usual number of purging) or nature of stool (more water than faecal matter). For age two months and over: 3 or more loose or liquid stools within 24 hours or 3 loose/liquid stools or fewer causing dehydration in the last 24 hours; enrolment per day was two patients with diarrhoea aged less than 5 years old and 2 patients aged 5 years or older; if the target number of patients in a particular age group was not met, the study overenrolled in the other group to meet the target of 4 patients per day during workdays. There were no exclusions (mHDM or NCS) based on prior reported antibiotic exposure.

### Bacterial culture

Stool samples were collected at hospital admission. Aliquots for transport and subsequent culture were stabbed into Cary-Blair transport media. The culture was performed via standard methods with both Thiosulfate–citrate–bile salts–sucrose agar (TCBS) and taurocholate-tellurite gelatine (TTGA) agar (16). Suspected *V. cholerae* colonies were serotyped with antibodies specific to O1 serogroup (Inaba and Ogawa) and O139 serogroup (16). Isolates were stored in glycerol at −80°C at the icddr,b, shipped on dry ice to the University of Florida, and re-isolated on Luria-Bertani (LB) agarose plates for nucleic acid analysis and storage at −80°C in glycerol.

### DNA extraction and sequencing

DNA extraction was performed on one pure colony grown in 5 ml of LB broth at 37°C at 220 rpm. DNA was extracted using the DNeasy Blood & Tissues kit following the manufacturer’s instructions (QIAGEN) followed by quantification using a NanoDrop spectrophotometer (Thermo Fisher Scientific) and Qubit fluorometer (Thermo Fisher Scientific). For each sample, approximately 200 ng was used to generate the sequencing library using the Nanopore Rapid Barcoding Kit 96 V14 (SQK-RBK114.96, Oxford Nanopore Technologies) following the instructions of the manufacturer. Briefly, the samples were made up to 10 µL with DNase-free water and 1 µL of rapid barcodes was added to each sample in groups of 96. The plates were incubated for 2 minutes at 30 °C followed by enzyme inactivation for 2 minutes at 80 °C. 2 µL per sample was used to pool the plates followed by concentrating using a 1:1 AMPure XP Beads clean up (Beckman Coulter). Beads were rinsed using two washes of 80% ethanol with 180-degree turns. The samples were eluted in 12 µL elution buffer EB (Oxford Nanopore Technologies). 11 µL of pooled library was ligated to 1 µL of pre-diluted Rapid Adapter and loaded on a PromethION R10.4.1 flow cell following priming (Oxford Nanopore Technologies). Whole genome sequencing was performed on a PromethION P2 Solo long-read sequencer with live base-calling enabled and Fast mode to monitor sequencing progress (Oxford Nanopore Technologies). The resulting Fast5 files were merged and converted to the POD5 format using Pod5 v.0.2.4 (Oxford Nanopore Technologies). The POD5 files were basecalled and demultiplexed per plate using guppy v.6.4.6 (Oxford Nanopore Technologies) using the *dna_r10.4.1_e8.2_400bps_sup* model with barcode and adapter trimming, and a minimum Q-score of 7. Four samples from the 277 collected failed to produce sufficient sequencing data and were removed from all downstream analyses.

### Phylogenetic analysis

Reads passing filtering for each demultiplexed sample were assembled using *Flye* v.2.9.1 using the option *-nano-hq* (17). The contigs were scaffolded for consistency using *RagTag* v.2.1.0 (18). To obtain a phylogeny using high-confidence single nucleotide variants (SNVs), reads were mapped to reference genome N16961 (RefSeq ID: GCF_900205735.1) using medaka v.1.11.3 (which employs minimap2 for mapping) (19–21). Short sequence reads of 776 publicly available 7PET genomes from around the world were additionally aligned to N16961 using BWA-MEM and variants were called using GATK (22,23). The resulting SNVs were used to construct a maximum-likelihood phylogenetic tree using RAxML v.8.2.12 with a bootstrap analysis over 500 replicates and with the GTRCAT model of nucleotide substitution, after removing indels and filtering out SNVs with QUAL scores lower than 40 (24). A 7PET isolate from 1971 Bangladesh (ERR025385) was used as the outgroup to root the tree (3). The assemblies were clustered using PopPUNK v.2.4.0, which compares the similarity of the whole genome *k-mer* set (25). The core genomes of these assemblies were also clustered using fastBAPS v.1.0.8 (26). The assemblies were screened for the presence of a set of known SXT-ICEs using BLASTN (10,27). An ICE was called as present if at least 90% of its sequence was covered in an isolate and the average identity of aligned sequences was larger than 95%. Ogawa or Inaba serotypes were inferred by comparing the *wbeT* sequences of the isolates - Inaba isolates contain mutations or structural variations that render the *wbeT* gene dysfunctional (28). The phylogenetic tree, combined with a heatmap showing the presence of ICEs and the geographical distribution were visualised in R using the package *ggtree* (29). Bayesian phylogenetic inference was performed on the BD-1.2 sub-lineage using BEAST to ascertain its transmission history, assuming a constant population size coalescence model with a strict clock rate and a γ distribution of base substitution heterogeneity (30).

### Pangenome analyses

Panaroo v.1.1.2 was used to estimate the pangenome of each *V. cholerae* lineage of interest (31). To measure the openness of pangenomes in BD-1 compared to BD-2, the model from Tettelin *et al*. using Heap’s law was fitted: *n*∼*N*^γ^, where *n* is the number of total genes and *N* is the number of genomes sampled (32). A higher γ value implies a higher rate of increase in gene numbers with an increase in genome numbers, hence a higher average divergence in genetic contents. Nei’s distance (π) was used to measure pairwise core distances, whereas Jaccard’s distance of the gene contents was used to measure the accessory distances. This was followed by plotting the distribution of accessory distances in BD-1 and BD-2 in a boxplot binned by core distances.

### Screening for known AMR genes

Known AMR genes in the Comprehensive Antibiotic Resistance Database (CARD) were screened in the *de novo* genome assemblies using CARD-RGI (33). To investigate deletions of AMR genes on ICEs, the ICE sequences were obtained by extracting the intervals between the two flanking genes, *prfC* (N16961_RS11085) and *gmr* (N16961_RS11090). Pairwise comparisons of ICE sequences were generated using BLAST and visualised with the Artemis Comparison Tool (ACT) (27,34).

### Screening for known phage-defence mechanisms

The presence of known phage-defence mechanisms in the *de novo* genome assemblies was inferred using DefenseFinder v.1.3.0 (35).

### Data availability

All DNA sequence data generated in this project is available in NCBI under BioProject PRJNA1174068.

### Statistical analyses and visualisation

Statistical analyses and visualisation were done using R. The R code used for plotting the figures is available from https://github.com/Chuhan-Qin/Vibrio_2024.

## Results

### Sample collection

A total of 277 *V. cholerae* isolates were obtained from stool samples collected as part of the nationwide study (15). One isolate was taken from each patient. We previously reported shotgun metagenomic sequencing from the same set of patients, who varied in dehydration severity from mild to severe (9).

### Long-read sequencing yields mostly complete genome assemblies

We successfully sequenced genomes from 273 of the 277 isolates using Nanopore technology (**Methods**). The median read N50 across samples was 6,598 bp, with a maximum of 43,509 bp. The reads were assembled into genomes ranging from 4.0 to 4.1 Mbp, consistent with the known range of *V. cholerae* genome sizes. Of note, the assembly sizes clustered around either 4.0 Mbp or 4.1 Mbp with very few in between (**Supplementary Fig. 1**). Most genomes (180/273; 65%) were assembled into two circular contigs, corresponding to the two complete chromosomes of *V. cholerae*. All 273 genomes were assembled into fewer than seven contigs, a marked improvement over the average of 73 contigs per genome in a recent study using short-read Illumina sequencing (36). Similarly, *V. cholerae* assemblies in the ENA database contained a median of 92 contigs, and only 202 of the 9,950 genomes (2.06%) were assembled into two circular contigs (accessed on 30 August 2024). In addition to the two circular chromosomes, we found only one other circular contig of 120 Kbp in isolate EN-1738, which was identified as an ICP1 phage.

### Major *V. cholerae* lineages identified in the phylogeny

To place our genomes in the context of known *V. cholerae* diversity, 776 publicly available *V. cholerae* 7PET genomes were downloaded from ENA and aligned with our genomes to construct a maximum-likelihood phylogeny (**Supplementary Table 1**). The tree was rooted with ERR025385, a 7PET isolate sampled from Bangladesh in 1971 (3). We found a median of 22 pairwise SNV differences amongst the 273 newly sequenced genomes, consistent with relatively little genetic diversity within 7PET. Most of these genomes fall within two lineages that have been co-circulating in Bangladesh since the 1990s, BD-1 and BD-2 (**Fig. 1a**) (37). All BD-1 isolates sequenced from our 2018-19 sampling were of the Ogawa serotype; this was before a serotype switch to Inaba in 2020 (7). All 2018-19 BD-2 isolates had an Inaba serotype.

**Fig. 1:**
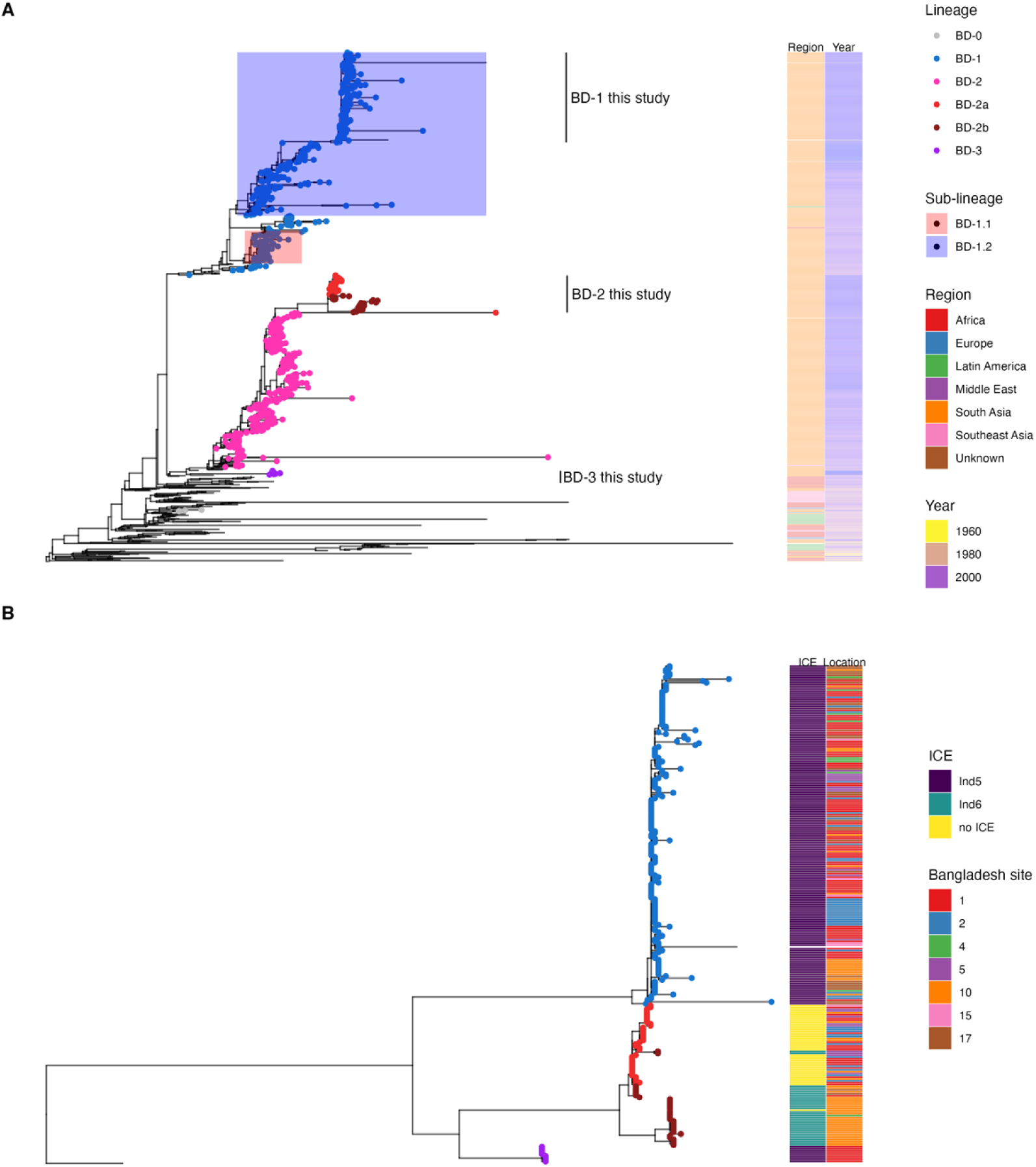
Major phylogenetic lineages of *V. cholerae* sampled across Bangladesh. **(A) Phylogenetic tree of newly sequenced *V. cholerae* genomes in the context of 776 publicly available genomes.** The maximum likelihood tree was rooted with a 1971 Bangladesh 7PET isolate as the outgroup (ERR025385). Leaves are coloured according to phylogenetic lineages: blue for BD-1, different shades of red for sub-lineages of BD-2, and purple for BD-3. Sub-lineages of BD-1 corresponding to BD-1.1 and BD-1.2, as defined in previous studies, were shaded. The heatmap shows the geographical regions and the year of sampling. **(B) Association between major lineages, ICE types, and sampling sites within Bangladesh.** The same outgroup ERR025385 and leaf colouring scheme were used as in panel (A). Other than the outgroup, only the 273 genomes sequenced in this study are included. The heatmap shows the presence of SXT-ICEs and the locations from which isolates were sampled within Bangladesh.

We also identified nine genomes forming a distinct monophyletic group (henceforth referred to as BD-3), which branches near the root of BD-2 (**Fig. 1a**). The publicly available genomes genetically closest to BD-3 are BD-2 isolates from the early 2000s, suggesting that BD-2 and BD-3 shared a common ancestor around that time. These lineages, based on SNVs in the core genome, showed a high concordance with the whole-genome *k-mer* clustering using PopPUNK (**Methods**). PopPUNK genomic cluster 1 (GC-1) corresponded to BD-1 (coloured blue in **Fig. 1**), whereas BD-2 was split into GC-2 and GC-3 (coloured shades of red in **Fig. 1**, henceforth referred to as BD-2a, BD-2b).

To support the phylogenetic relationships among *V. cholerae* lineages at a higher resolution, a second maximum-likelihood tree was constructed using the 273 new genomes only (**Fig. 1B**). Both trees identified BD-1, BD-2, and BD-3 as distinct monophyletic groups with bootstrap support of 100%. Consistent with our earlier metagenomic sequencing, two types of ICEs, *Vch*Ind5 and *Vch*Ind6, are found in the isolate genomes (9). All BD-2a isolates lacked an SXT-ICE, consistent with the shorter median length of BD-2a assemblies, which was 91,825 bp shorter than that of BD-2b (Welch two sample *t*-test: *p* < 2.2*10^-16^), close to the average size *Vch*Ind6 at 94,599 bps (**Supplementary Fig. 1**). Other than occasional ICE gain or loss events, the ICE type is generally a stable feature within each lineage (**Fig. 1B**), with BD-1 and BD-3 containing *Vch*Ind5 and most BD-2b genomes containing *Vch*Ind6.

### Geographical and temporal distribution of *V. cholerae* genomic diversity

To quantify the genomic diversity of *V. cholerae* between time points and locations, we compared the Shannon diversity of PopPUNK genomic cluster (GC) composition. The GCs largely map to phylogenetic lineages: GC-1 to BD-1, GC-2 to BD-2a, GC-3 to BD-2b, and GC-4 to BD-3 (**Supplementary Fig. 2**). However, in addition to the core genome SNVs used to construct the phylogenetic trees, PopPUNK also uses accessory genome content, resulting in higher genetic resolution: 39 GCs compared to only 4 major phylogenetic lineages. For example, a genome with distinct gene content would be classified as a separate GC, despite branching within BD-1 or BD-2 (**Supplementary Fig. 2**). Such gene content ‘outliers’ tend to be rare, and are considered together as ‘others’ for visualisation purposes but counted separately to calculate Shannon diversity (**Fig. 2**).

**Fig. 2:**
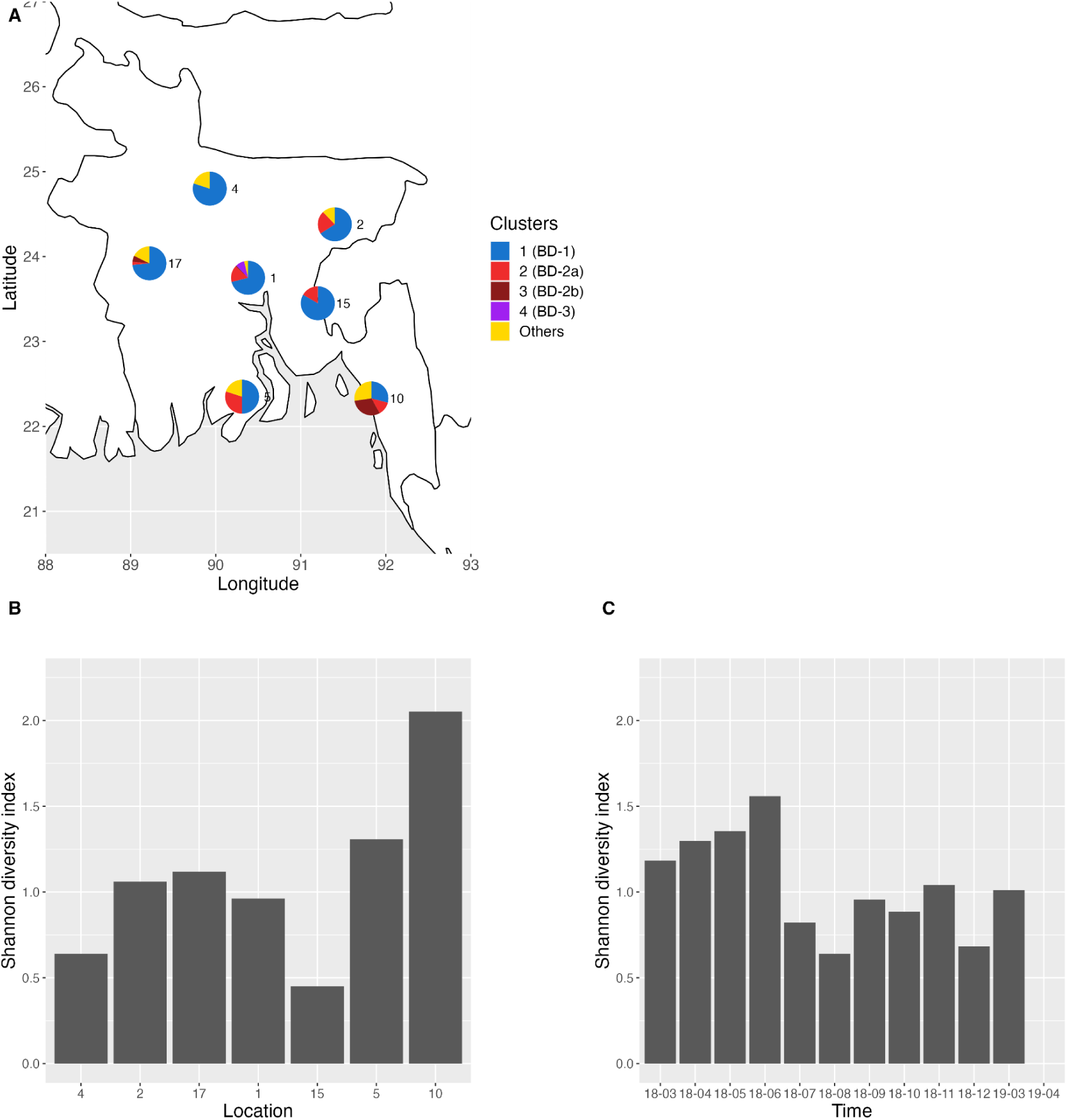
**(A) Geographical distribution of *V. cholerae* genomic clusters across Bangladesh.** Isolates are coloured by PopPUNK genomic clusters, matching with lineages coloured as in Fig 1. ‘Others’ denotes rare clusters only observed once. **(B) Shannon diversity of genomic clusters by location.** Locations are sorted by latitude from the north to the south (left to right). **(C) Shannon diversity of genomic clusters by month.** The *X*-axis shows the sampling period in chronological order from March 2018 (18–03) to April 2019 (19–04). Note that the final time point was only sampled from Dhaka, and only contained BD-3, resulting in a Shannon diversity value of zero.

The distribution of *V. cholerae* GCs varied among sampling locations across Bangladesh (**Fig. 2A, B**). The sites in the north, close to the Bangladesh-India border, contained predominantly BD-1.2 (GC-1, >70%). By contrast, we observed a higher diversity of lineages in the two southern sites on the coast of the Bay of Bengal, marked by a mixture of BD-1 (GC-1) and BD-2 (GC-2 and GC-3). Previous work inferred an Indian origin for BD-1.2 in Bangladesh, which is consistent with the enrichment of BD-1.2 in parts of Bangladesh physically closer to India (37). Further supporting an Indian origin, all BD-1 isolates in this study clustered with publicly available BD-1.2 genomes, and were genetically closer to Indian BD-1 isolates from the early 2010s than to Bangladeshi isolates of the same time **(Fig. 1A)**. To more formally infer the divergence time and geographic source of BD-1.2, we used Bayesian ancestral reconstruction to infer the most recent common ancestor of BD-1.2 (**Methods**). We estimated the most recent common ancestor of BD-1.2 in 2002 (95% HPD interval: 2000 - 2003) and was of Indian origin. Site 10, which is located near a refugee camp at Cox’s Bazar, was also distinct in its lineage composition, containing a significant number of both BD-2a (GC-2) and BD-2b (GC-3) and high Shannon diversity (**Fig. 2B**), whereas most other sites contained only one or the other (38). BD-3 (GC-4) was found only in Dhaka.

Lineage diversity varied less over time (s.d.=0.41; **Fig 2C**) than over sampling locations (s.d.=0.52, **Fig. 2B**). The 2019 samples all came from Dhaka, and samples from April 2019 had a Shannon diversity of zero as all isolates from that month were BD-3. When excluding the 2019 samples, the standard deviation of Shannon diversity over time is only 0.30, much less than the variation over space. A shift from BD-2 to BD-1 has been previously reported in Dhaka from 2018 to 2019 (7). We also observe a higher frequency of BD-1, particularly at later time points during our sampling period (**Supplementary Fig. 3**), suggesting that the shift toward BD-1 may be a nationwide phenomenon. When examining the wider collection of publicly available genomes (**Supplementary Table 1**), we found that shifts in the dominant lineage had occurred previously in Bangladesh - from predominantly BD-1 in 2010 (90.9%) to a mixture of BD-1 and BD-2 in 2011-2012, followed by a predominantly BD-2 population (96.7%) between 2013 and 2017. Therefore, the alternating dominance of BD-1 and BD-2 could be explained by fluctuating or frequency-dependent selection rather than a directional lineage replacement.

### Lineage BD-2 has a more open pangenome than BD-1

Next, we compared the accessory gene content of BD-1 and BD-2. A pangenome is defined here as the entire set of genes from all isolates within a lineage. We found that BD-2 had a more ‘open’ pangenome (Tettelin’s γ = 0.0291), with more genes added to the pangenome per sequenced genome compared to BD-1 (γ = 0.0172; **Fig. 3A**). Pairwise Jaccard distances between accessory gene content were consistently higher in BD-2, and this was true across all core genome distance bins (Welch two sample *t*-test: *p* < 2.2*10^-16^, **Fig. 3B**). This implies that the differences in pangenome variation are not due to differences in phylogenetic depth between BD-1 and BD-2. The high level of pangenome variation within BD-2 is likely due to its subdivision in BD-2a, which lacks an ICE, and BD-2b, which contains the *Vch*Ind6 ICE.

**Fig. 3:**
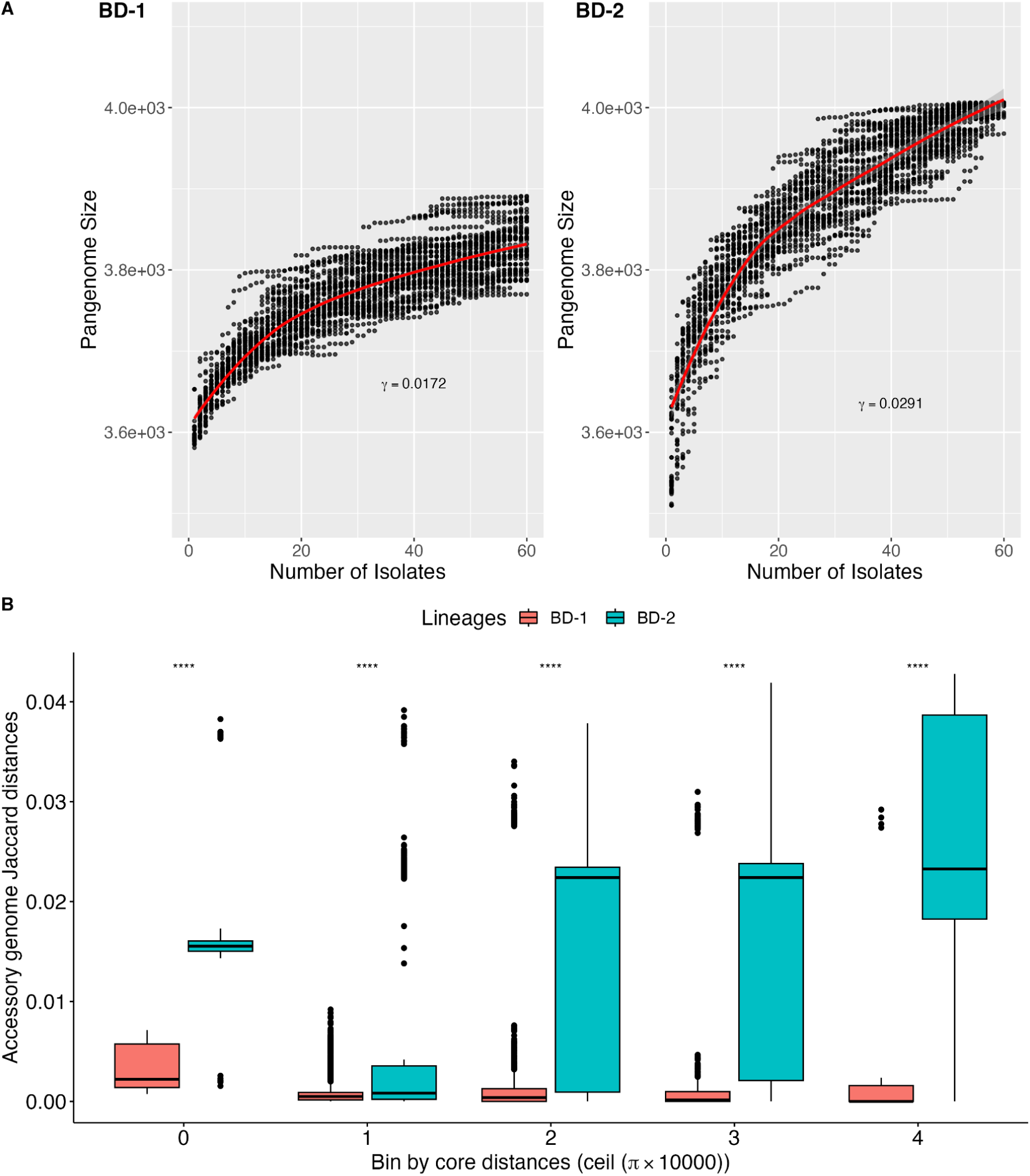
BD-2 encodes a more diverse ‘open’ pangenome than BD-1. **(A) Pangenome size grows faster with sampling effort in BD-2 than BD-1.** The *X*-axis shows the number of isolates in each sub-sample. The *Y*-axis shows its corresponding pangenome size (number of genes). For each lineage, 50 subsets were sampled (black points). The red line shows the median. γ is the index in Tettelin’s model: *n*∼*N*^γ^; a higher value indicates a more open pangenome **(B) Distribution of within-lineage accessory distances in different core genome distance bins.** The bins were set according to the core genome Nei’s distances (π). The boxplots show the range of the within-lineage accessory genome Jaccard distances. Boxes were coloured in either red (BD-1) or green (BD-2). ‘****’ denotes a *p*-value < 2.2*10^-16^ using the *t*-test.

### Variation in AMR gene profiles

We used the CARD database to identify annotated AMR genes in our sample of genomes. We found 16 known AMR genes present in one or more genomes using the strict matching criteria (**Supplementary Table 2**). The major lineages had distinct presence/absence profiles of these 16 genes (**Fig. 4A**). Multidrug efflux pump *rsmA* (for fluoroquinolone, diaminopyrimidine, and chloramphenicol) was found in all 273 genomes. So was CRP, a global regulator for the efflux pump MdtEF, which removes fluoroquinolone, macrolide, and beta-lactam from bacterial cells (39,40). Note that the screening for CRP did not consider a point mutation that was shown to render it dysfunctional (40). The loss of the SXT-ICE element in BD-2a profoundly affected its AMR gene profile: almost all genomes other than BD-2a contain APH(6)-Id or APH(3)-Ib (>99%), a phosphotransferase of aminoglycoside, which is completely absent in BD-2a (**Fig. 4A**). The same pattern was observed for the sulfonamide and trimethoprim resistance genes, *dfrA* and *sul2*, both of which were absent in BD-2a but had high frequencies in the other lineages. AMR genes against vancomycin (*vanY* and *vanT*); carbapenem (*varG*); fluoroquinolone (*parE*); and colistin (*almEFG*) were also prevalent across lineages (>95%). A point mutation involved in chloramphenicol inactivation *catB9* was present in all but one BD-1 isolate. The chloramphenicol efflux pump *floR* is a hallmark of *Vch*Ind5; it was absent in BD-2 but present in almost all BD-1 and BD-3 isolates. Consistent with previous findings, *tetA*, an efflux pump for tetracycline, is located on the *Vch*Ind6 and is present in all BD-2b but none of the other genomes (13).

**Fig. 4:**
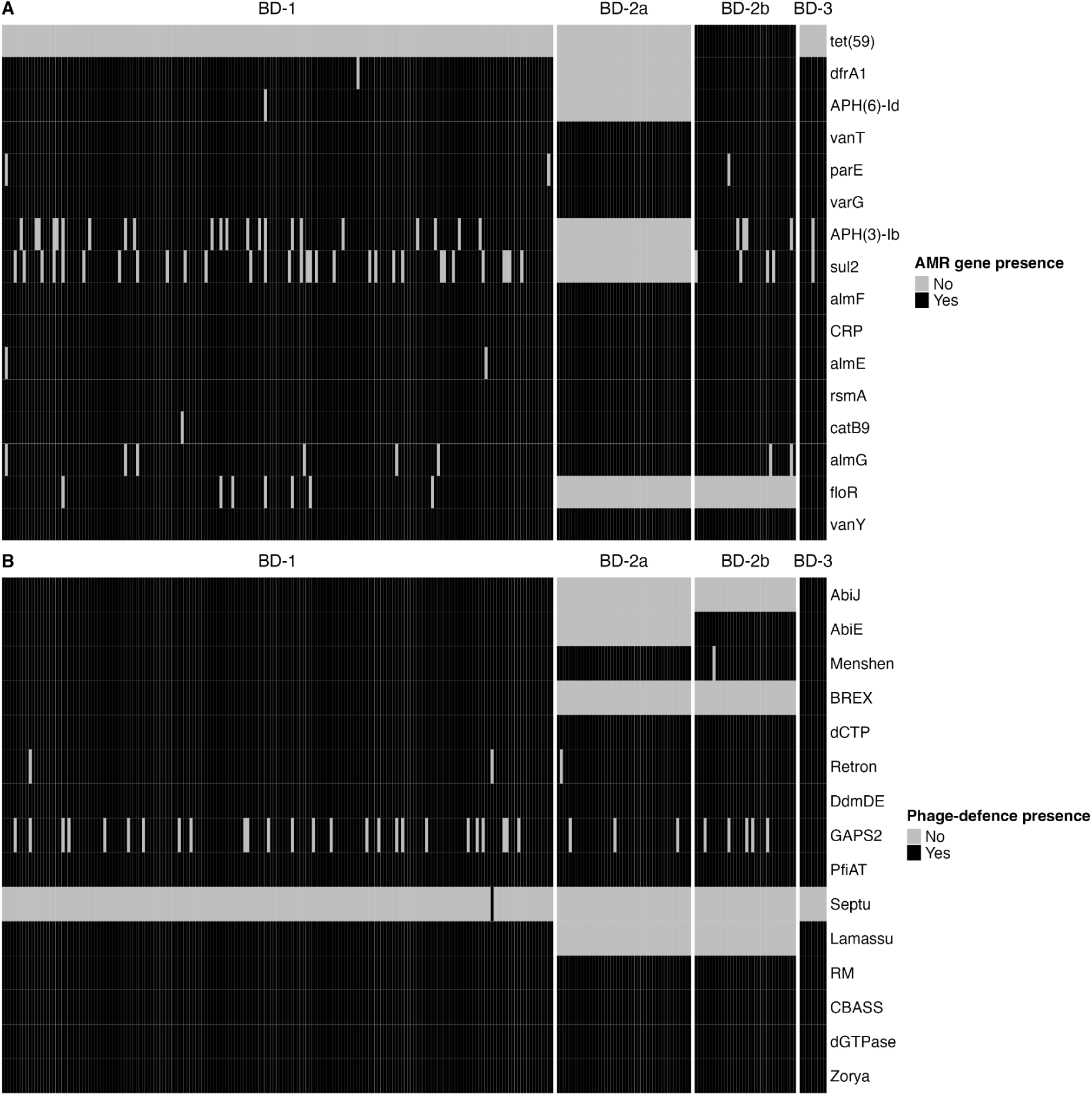
Distribution of AMR and phage-defence genes vary across major lineages of *V. cholerae*. The *X*-axis shows the isolates and is sorted by lineages. (**A**) The *Y*-axis shows the 16 AMR genes from the CARD that occurred at least once in the genome assemblies (33). (**B**) The *Y-*axis shows the 15 phage-defence genes identified by DefenseFinder in the genome assemblies (35). For both panels, black represents the presence whereas grey indicates the absence.

Notably, two AMR genes on the SXT-R391 ICE (*qnr_Vc_* and *mphA*) that were previously associated with ciprofloxacin and azithromycin resistance in a 2006 sample of *V. cholerae* genomes from Bangladesh were absent in the 2018-2019 genomes(13). To compare the structure of the SXT-R391 ICEs that harboured these genes, their sequences were obtained by extracting bases between the genes *prfC* and *gmr*, flanking the ICE insertion site. A *Vch*Ind6 from a 2006 BD-2 isolate (EN-1443) that contains *qnr_Vc_* and *mphA* was compared to another *Vch*Ind6 (EN-1581) from a 2018 BD-2 isolate (**Fig. 5A**). All *Vch*Ind6 from 2018 share this loss of *qnr_Vc_* and *mphA*. A 2018 *Vch*Ind5 ICE from a 2018 genome also contained this deletion relative to the ICE from 2006 (**Fig. 5B**). Thus, *qnr_Vc_* and *mphA* were absent in the 2018-2019 genomes due to one or more deletion events of an ICE region that was present in 2006 (13). It is unclear if these genes were deleted due to fitness costs, if they were replaced with more effective resistance mechanisms elsewhere in the genome (since ciprofloxacin and azithromycin are still commonly used antibiotics for cholera patients in Bangladesh), or a combination of these factors.

**Fig. 5:**
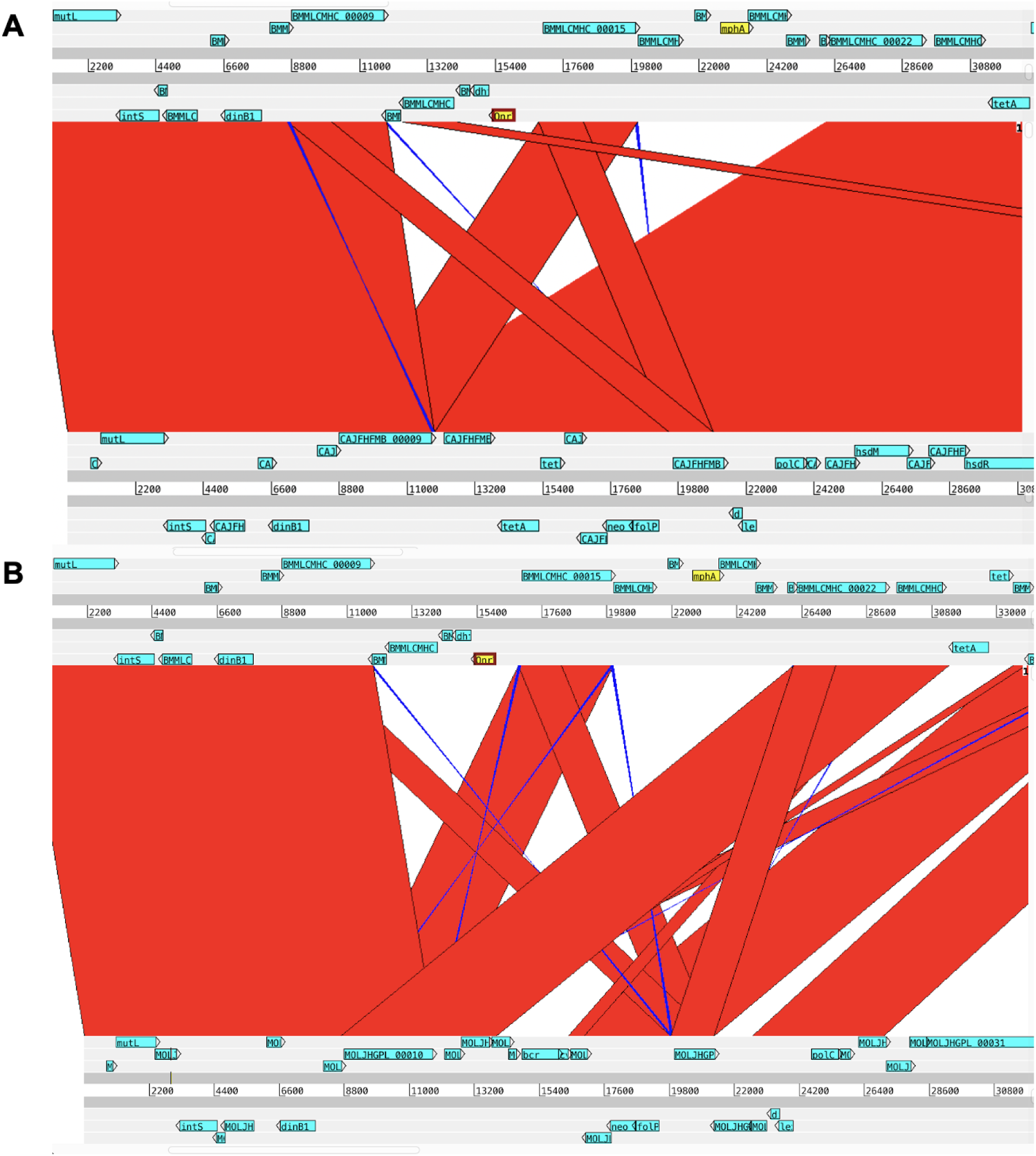
Local genome comparison plot of the ICE region containing AMR genes lost between 2006 and 2018. (A) *Vch*Ind6 from a 2006 BD-2 isolate (EN-1443) that contains *qnr_Vc_* and *mphA* compared to *Vch*Ind6 from a 2018 BD-2 isolate (EN-1581). (B) *Vch*Ind6 from EN-1443 compared to *Vch*Ind5 from another 2018 BD-2 isolate (EN-1469). Red blocks show aligned regions in the same order, whereas blue blocks show reversed aligned regions. Two important AMR genes absent from the 2018 genomes are coloured yellow.

### Variation in phage-defence gene profiles

We next identified known phage defence genes in our genomes using DefenseFinder (35). Despite being relatively distantly related (**Fig. 1**), BD-1 and BD-3 genomes all contain 13 or 14 of the 15 identified defence systems (**Fig. 4B**). This suggests that defence systems such as *Abi*, BREX, and Lamassu were present in the common ancestor of BD-1, 2 and 3, and were lost in BD-2. Other phage defence systems such as DdmDE, Zorya and restriction-modification systems are universal to all sequenced genomes (**Fig. 4B**). The Lamassu system, commonly referred to as *DdmABC* in the *Vibrio* literature, acts in conjunction with *Abi* to kill phage-infected bacterial cells before phage replication can be completed, thereby providing population-level immunity (41). BD-1 and BD-2 encode different variants of VSP-II, where *DdmABC* is encoded, likely explaining the lineage-specific differences (38). Consistent with a prior study linking the BREX defence system to *Vch*Ind5, BREX genes were present in all *Vch*Ind5-containing genomes (BD-1 and BD-3) but none of the BD-2 isolates (10)(**Fig. 4B**). Together, these results show that co-circulating lineages have distinct profiles of both antibiotic and phage resistance genes.

## Discussion

In this study, we analysed *V. cholerae* genomes from cholera patients across Bangladesh over one year to assess genetic diversity using long-read sequencing. This revealed substantial diversity of different *V. cholerae* lineages encoding distinct profiles of AMR and phage resistance genes co-circulating across the country. Although the sample in this study covered a wide geographical region across Bangladesh, the sample size per month was still rather small, and from early 2019 onward, isolates were exclusively sourced from icddr,b. Despite these limitations, we identified an intriguing north-south gradient of genomic diversity, with a dominant lineage likely introduced from India to the north, and greater diversity persisting in the south. This pattern merits further investigations in larger and longer sampling efforts.

We also demonstrate the utility of long-read nanopore sequencing to resolve structure variation in *V. cholerae* genomes and to infer phylogenetic relationships. The numbers of single nucleotide variants in the long-read genomes were comparable to those in short-read genomes over similar spatial and temporal scales, suggesting that sequencing errors are not excessive in the long-reads after quality filtering (42). The benefit of long-read sequencing technology is that large-scale genomic structures can be resolved at higher accuracy – for instance, we observed the deletion of SXT-ICE in a sub-lineage. Of the 273 genomes sequenced, 65% were assembled into two circular contigs, corresponding to the two chromosomes in *V. cholerae*, with the remaining assemblies failing to yield closed chromosomes. With DNA extraction protocols optimised for long reads, the completeness of assemblies could be further improved in future analyses.

Two distinct *V. cholerae* lineages, BD-1 and BD-2, have been predominant in Bangladesh since the late 1990s (8). Between 1999 and 2017, BD-2 increased in frequency compared to BD-1 (8), until the BD-1.2 sub-lineage emerged as the cause of a major outbreak in 2022. Filling the gap between 2017 and 2022, our analysis shows that BD-1.2 dominated BD.2 throughout 2018, particularly in the north of Bangladesh. However, BD.2 persisted through the end of our sampling in 2019. Our phylodynamic and geographic analyses support an Indian origin of BD1.2, which likely arrived in Bangladesh from the north. Furthermore, we report a new lineage, BD-3, which only appeared in our samples from the icddr,b sampling site in Dhaka. This lineage appeared throughout the sampling period but became particularly frequent in 2019, when it comprised the majority of genomes at icddr,b. BD-3 has a similar AMR and phage resistance gene profile as BD-1, suggesting that it could compete with BD-1 in the future. Ongoing monitoring will be required to determine if this occurs.

Monir *et al.,* suggested serotype switching as a possible explanation for the success of BD-1.2 over BD.2 (7). Although we observed higher proportions of Ogawa serotype towards the end of our sampling period in early 2019, some Inaba isolates persisted. The O-antigen may be under negative-frequency dependent selection, such that no serotype reaches fixation and diversity is maintained (43). Indeed, Ogawa and Inaba serotypes have co-existed in O1 7PET since the beginning of the 7th pandemic in the 1960s, and serotype switchings in both directions have been frequently observed (28,44,45).

Negative frequency-dependent selection could also arise from the co-evolution of *V. cholerae* and its virulent phages. *V. cholerae* lineages differ in the types of ICE they encode, and some genomes lack an ICE entirely (10). Different ICE types encode different genes involved in both phage and antibiotic resistance (10). For example, in our study, both BD-3 and BD-1 contained *VchInd5*, whereas BD-2 was split into sub-lineages that either possessed *VchInd6* or lacked an ICE. Intriguingly, a significant proportion of the isolates lacked an SXT-ICE despite the potential benefits it confers, such as antibiotic and phage resistance (10,14). LeGault *et al.,* posited that it may be beneficial for a *V. cholerae* population to include multiple lineages with distinct ICEs to increase the diversity of defence mechanisms against the ICP1 phage(10). Strains without ICEs may be beneficial at the population level, as ICP1 released from ICE-free *V. cholerae* lack epigenetic modifications necessary to evade ICE-encoded phage-restriction systems, thereby reducing the overall burden of phage predation (9,10). Our data are consistent with the hypothesis that negative frequency-dependent selection for rare ICE types, or stabilising selection to maintain ICE-free cells at low frequencies, might explain the lack of a complete replacement of BD-2 by BD-1.2.

Antibiotic resistance has become widespread in clinical *V. cholerae*. A study of 443 *V. cholerae* isolates in the endemic region of India found that more than 99% of these were multidrug-resistant (46). Consistent with this observation, 16 known AMR genes were identified in our data set, conferring resistance to a wide variety of antibiotics - including aminoglycoside, sulfonamide, trimethoprim, vancomycin, carbapenem, fluoroquinolone, macrolide, colistin, chloramphenicol, and tetracycline. Each genome contained at least ten of these 16 genes. The acquisition of AMR in *V. cholerae* has likely occurred via mobile elements such as ICEs, which can disseminate AMR genes between genetically distant strains (46). In our dataset, distinct ICE variants are stably associated with distinct lineages, with occasional ICE gain or loss within a lineage. Notably, a variable ICE region containing two genes strongly associated with ciprofloxacin and azithromycin resistance in 2006, *qnr_Vc_* and *mphA*, were completely absent in the 2018-2019 genomes. Extensive gene conversions have been observed between SXT-ICEs, which could provide a mechanism for these deletions (14). The absence of these canonical ciprofloxacin and azithromycin resistance genes would necessitate alternative mechanisms if *V. cholerae* strains were to remain resistant to both drugs, as they likely do (47,48). Further study will be needed to determine if contemporary *V. cholerae* remain resistant to these drugs, and if so, by which genetic mechanisms.

In summary, our results highlight the genetic diversity present in clinical *V. cholerae* across Bangladesh over a one-year outbreak season. In contrast to the lineage replacement events that appear to take place over decades in subsequent waves of globally successful clones, we observe that multiple distinct lineages, with distinct gene content, coexist within a year on a nationwide scale (3). The north-south gradient of genomic diversity suggests geographically structured environmental pressures and transmission routes, which deserve further study.

## Acknowledgements

We thank the patients for participating in the studies from which the strains were obtained, and the research teams at the icddr,b who made this work possible. We are grateful to Brittney Johnson, Randy Autrey, and Krista Berquist for their administrative expertise at University of Florida Department of Pediatrics.

## Supplementary materials

**Supplementary Fig. 1:**
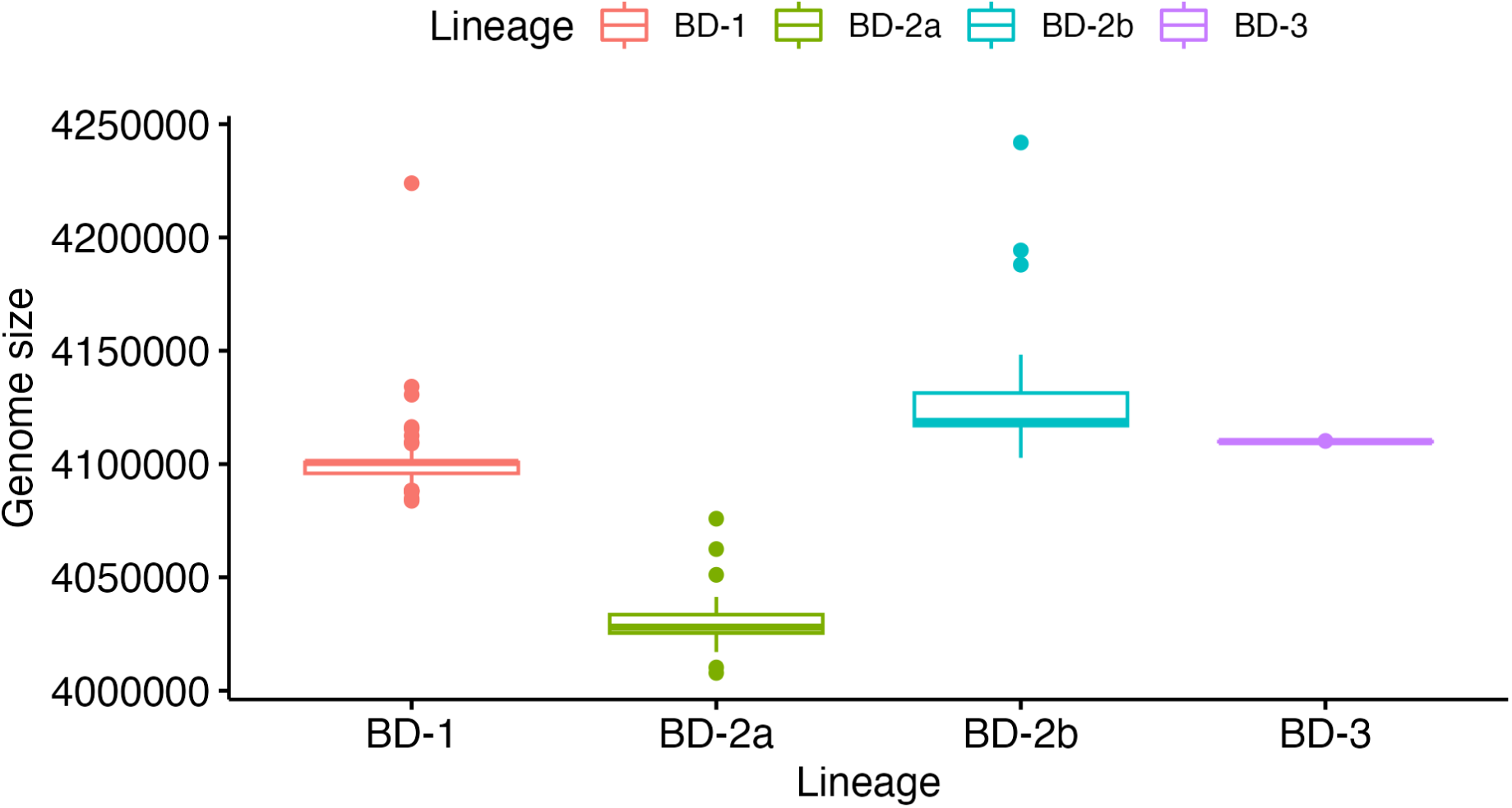
Distribution of genome assembly length by *V. cholerae* lineage. Boxplots showing the genome size distribution on the *Y*-axis which are grouped by lineages along the *Y*-axis. Lineages are also indicated by different colours.

**Supplementary Fig. 2:**
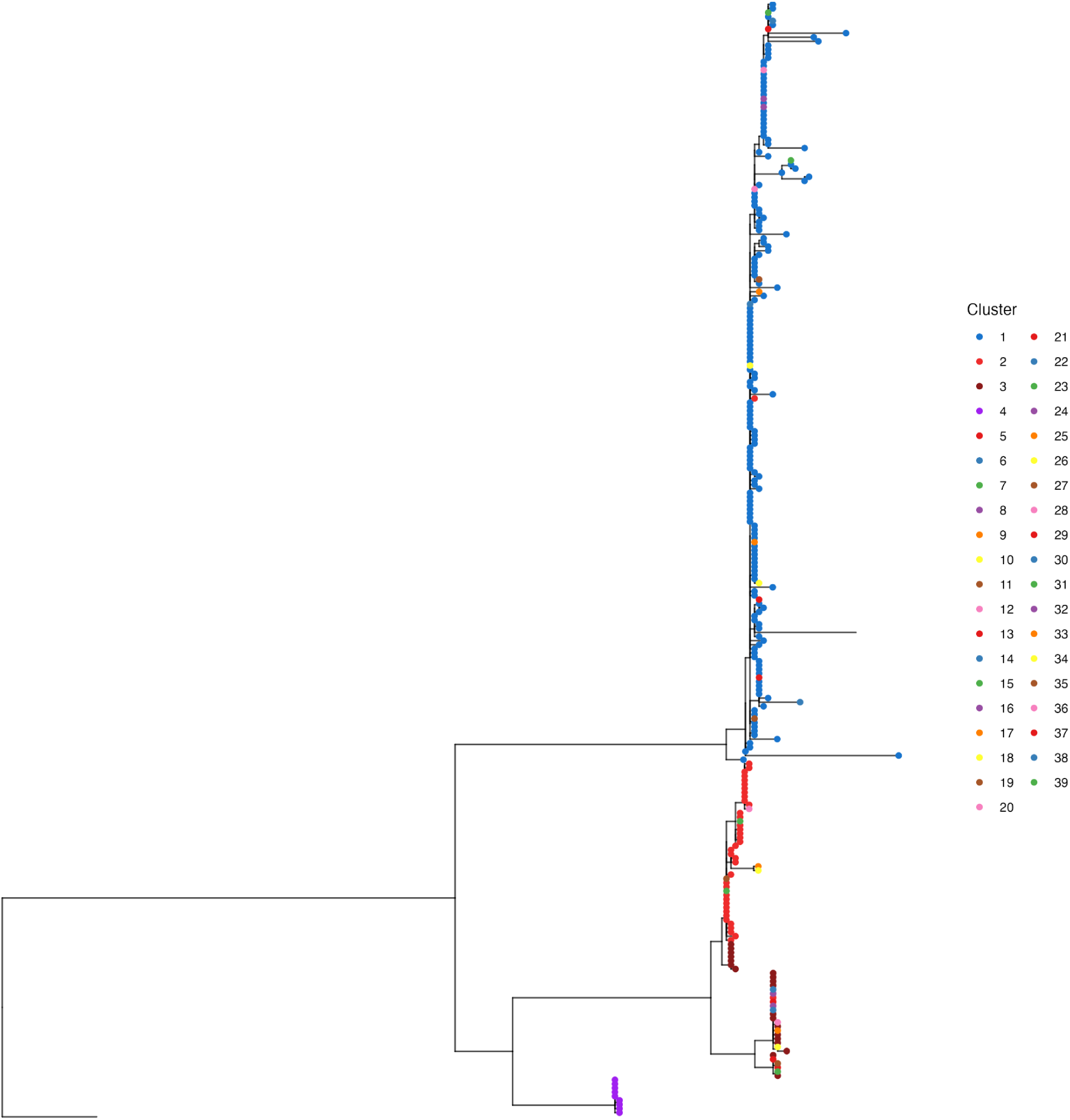
PopPUNK clustering of newly-sequenced *V. cholerae* genomes. The maximum likelihood tree was rooted with a 1971 Bangladesh 7PET isolate as the outgroup (ERR025385). Leaves are coloured according to PopPUNK clusters, Clusters 1 to 4 were coloured using the same scale as the corresponding BD lineages in Fig. 1. Clusters 5 to 39 are singletons that mostly cluster within BD-1, BD-2a or BD-2b, but contain distinct accessory genome contents and are shown in different colours.

**Supplementary Fig. 3:**
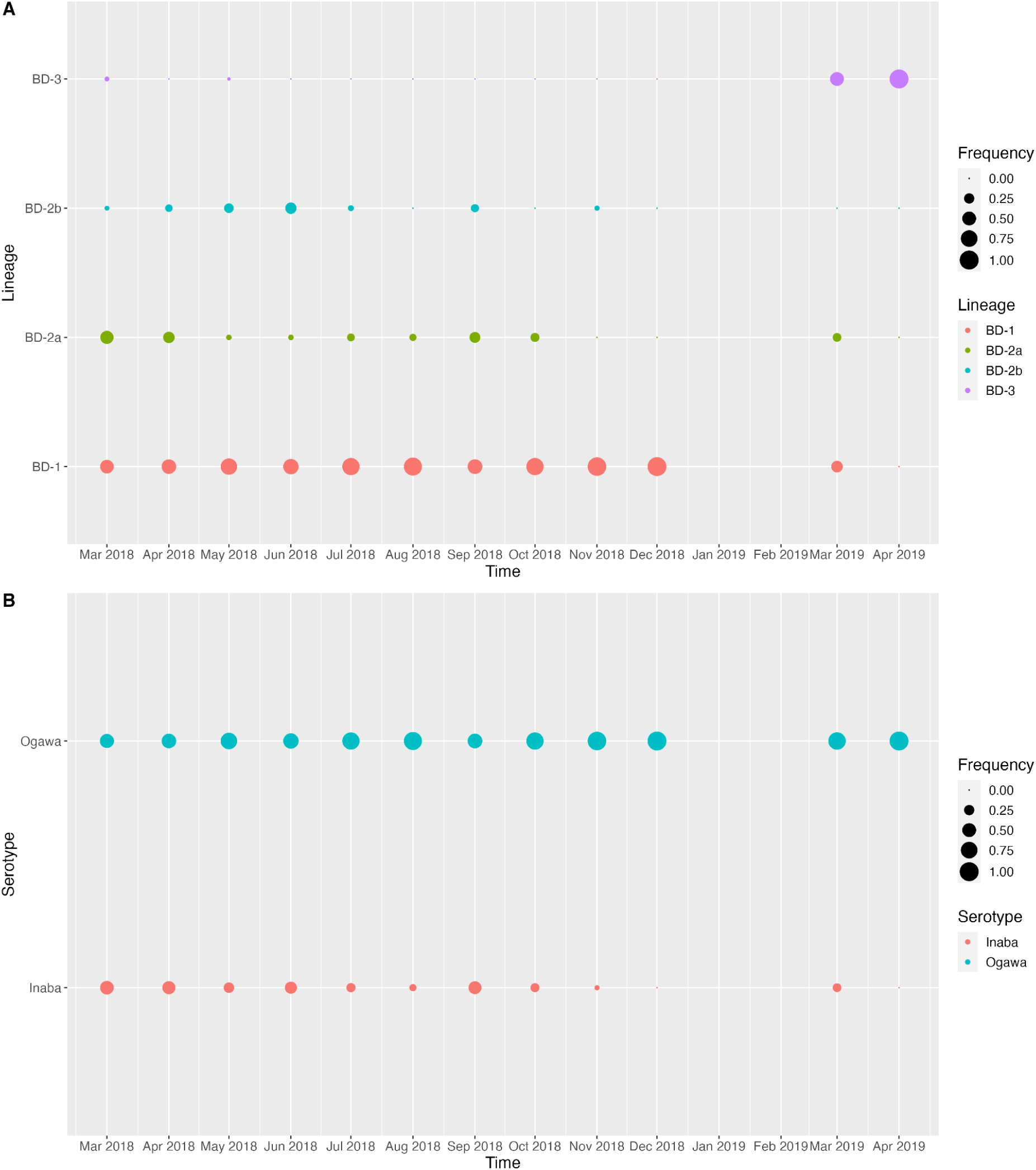
(A) Frequency of phylogenetic lineages over time. *X*-axis shows the time of sampling; *Y*-axis shows the lineages in different colours. The size of the dots indicates the proportion of each lineage within isolates from the same month. Note that the isolates in 2019 were from icddr,b only. **(B) Frequency of serotypes over time.** *X*-axis shows the time of sampling; *Y*-axis shows the Ogawa and Inaba serotypes in different colours. The size of the dots indicates the proportion of each serotype within isolates from the same month.

**Supplementary Table 1:** Metadata of newly sequenced and publicly available genomes used to construct the phylogeny in **Fig. 1**. Columns contain information on the year of isolation, the country of origin, the data set, and the lineage.

**Supplementary Table 2:** The first sheet shows the presence of AMR hits from the CARD-RGI output, ‘1’ denotes presence and ‘0’ denotes absence. The second sheet shows the annotation for each AMR hit, including information on frequency in the data set, the drug class and antibiotics affected, and the resistance mechanism.

